# Genetic screen identifies cell wall enzyme is key for freshwater survival of *Francisella tularensis*

**DOI:** 10.1101/2024.11.21.624769

**Authors:** Aisling Macaraeg, Hannah S. Trautmann, Kathryn M. Ramsey

**Affiliations:** Department of Cell and Molecular Biology, University of Rhode Island, Kingston, Rhode Island, USA

## Abstract

Human infection with *Francisella tularensis*, a potentially lethal bacterial pathogen, typically occurs after exposure to contaminated water, soil, food, or an infected animal. While *F. tularensis* can persist in environmental sources over long periods of time, the genetic requirements that permit its long-term viability are not understood. To address this question, we developed a laboratory model for persistence of *F. tularensis* in fresh water, finding that viable cells could be recovered for 3 – 8 weeks after incubation at 4°C. Using this model, we took an unbiased, transposon insertion sequencing approach to identify genes critical for this persistence of *F. tularensis* cells. We found that mutants in *mpl*, a gene encoding murein peptide ligase, are defective for persistence in fresh water. Previous studies had identified *mpl* as critical for intramacrophage survival. Murein peptide ligase plays a role in peptidoglycan recycling, suggesting that *F. tularensis* uses this enzyme to maintain cell wall integrity during hypoosmotic and intramacrophage stress conditions. Our results highlight the importance of understanding how bacterial cell envelopes have evolved and adapted to maintain their integrity in a variety of stress conditions.

## Introduction

*Francisella tularensis* is a Gram-negative facultative intracellular pathogenic bacterium and the causative agent of the disease tularemia. While *F. tularensis* is not directly transmitted between humans, a wide variety of environmental exposures can lead to infection, including arthropod bites, handling infected animals, eating contaminated food, and drinking contaminated water (Sjöstedt, 2007). Environmental persistence of *F. tularensis*, particularly in water sources, can lead directly to human infection as well as amplification in local animal populations and subsequent outbreaks in humans (Hennebique et al., 2019).

The presence and persistence of *F. tularensis* in a variety of environmental sources, and particularly aquatic environments, has been well-described (reviewed in Hennebique et al., 2019). Since the 1950s, it has been recognized that water and mud contaminated with *F. tularensis* can remain infectious for as long as 16 months (Parker et al., 1951). Subsequently, many studies have examined the survival of *Francisella* species in water under varying environmental conditions such as salinity, temperature, and presence of predators or possible hosts (Forsman et al., 2000; Thelaus et al., 2009; Berrada et al., 2011; Gilbert and Rose, 2012; Siebert et al., 2020; Hennebique et al., 2021; Golovliov et al., 2021). A gene encoding the mechanosensitive channel FtMscS has been identified to be important for *F. tularensis* to survive the osmotic shock that would occur during the transition between mammalian hosts into fresh water (Williamson, 2017). Beyond this, however, very little is known about the genetic requirements for long-term survival of *F. tularensis* to survive in aquatic conditions.

Here, we took an unbiased, genome-wide screening approach to identify genes necessary for survival of *F. tularensis* subsp. *holarctica* LVS (the live vaccine strain) in fresh water. First, we defined conditions that led to long-term viability of *F. tularensis* LVS in fresh water. We then modified an existing transposon delivery vector (Ramsey, Ledvina, et al., 2020) to permit sequencing of transposon insertion libraries made using the InSeq protocol (Goodman et al., 2011) using cost-effective technology (MiSeq). Using this transposon delivery vector, we generated a transposon insertion library in *F. tularensis* LVS cells. Finally, we used transposon insertion sequencing (Tn-Seq) to assess the ability of these mutant cells to persist in fresh water. This led to the identification of a candidate gene necessary for long-term survival of *F. tularensis* in fresh water, *mpl*, which encodes murein peptide ligase. This work is the first to identify a specific *F. tularensis* gene important for persistence in aquatic environments and suggests that other genes, particularly those involved in cell wall maintenance, may also be important for cellular survival in this key environmental niche.

## Results

### A model system to examine viability of *F. tularensis* in fresh water

We first sought to establish conditions that would permit long-term persistence of viable *F. tularensis* in aquatic conditions. We chose to use water taken from a local river (Beaver River, RI, USA) and examined how varying temperatures would impact the survival of *F. tularensis* LVS in this water over time. In particular, we inoculated *F. tularensis* LVS into filter-sterilized fresh water held at three different temperatures: 4°C, 16°C, and 25°C, in triplicate (**Fig. 1**). Throughout the next weeks, these replicates were sampled at regular intervals to evaluate bacterial viability by dilution and enumeration of colony-forming units. We continued sampling and assessing viability until two weeks elapsed with no colonies detected from undiluted samples (**Fig. 2**). We found that at 4°C, *F. tularensis* LVS cells remained viable for the longest amount of time, through 35 days, with none detected after 42 days of incubation. In comparison, the last viable cells were detected after 14 days when fresh water cultures were incubated at 16°C and 7 days after incubation at 25°C.

**Figure 1.**
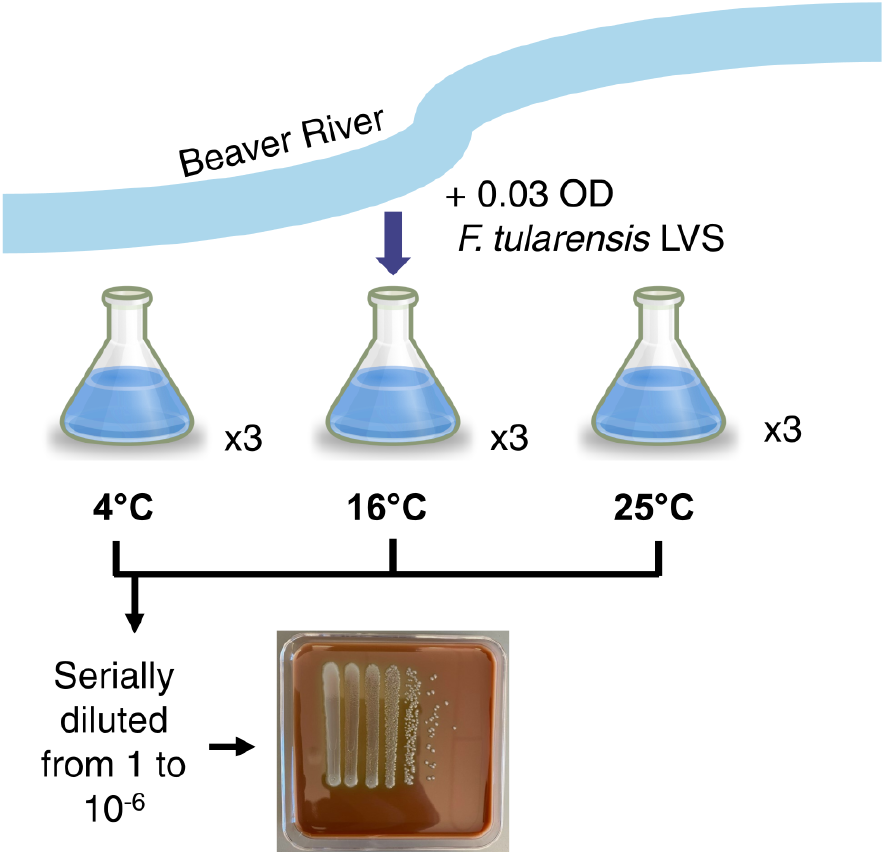
Workflow for development of fresh water persistence model. Water was retrieved from the Beaver River and was filter sterilized. *F. tularensis* LVS was diluted into filter-sterilized fresh water to an optical density of 0.03 and distributed into nine flasks. Flasks were incubated at 4°C, 16°C, or 25°C in triplicate. Each week, the flasks were sampled and the remaining viable *F. tularensis* cells were determined by dilution and enumeration of colony-forming units. A representative plate is shown.

**Figure 2.**
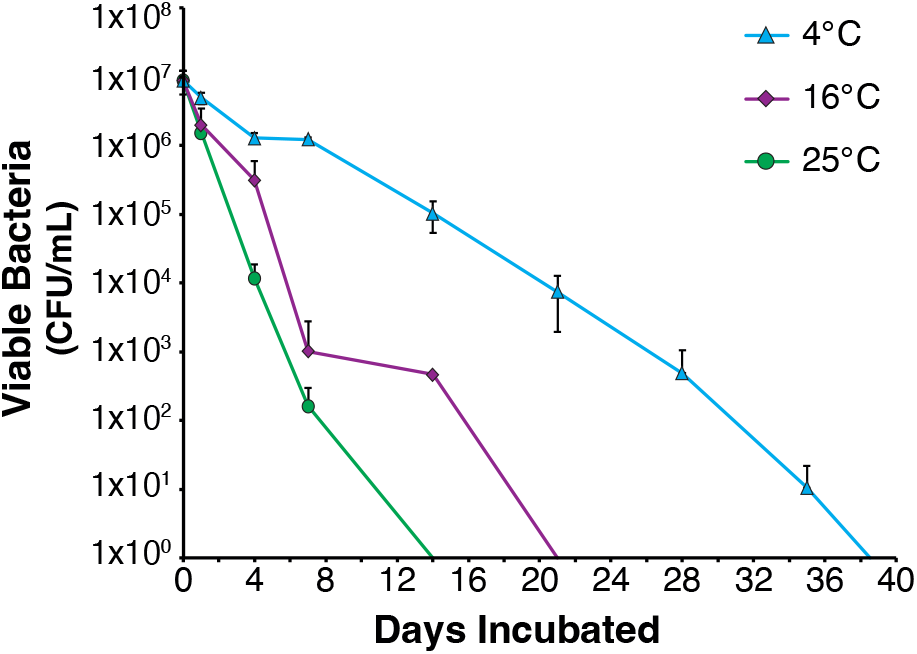
Survival of *F. tularensis* LVS in fresh water at different temperatures. Average colony forming units (CFU) per mL recovered at indicated time points. Cells incubated at 4°C are represented by blue triangles, cells incubated at 16°C are represented by purple diamonds, and cells incubated at 25°C are represented by green circles. Each point represents the average number of cells recovered from three independent flasks. Error bars represent 1 SD.

We sought to assess the reproducibility of these results by performing two more independent experiments to examine survival of *F. tularensis* LVS in fresh water at 4°C (**Fig. 3**). We found that the length of time *F. tularensis* LVS cells remained viable varied, from 21 to 56 days, but was consistently longer than cells incubated at 16°C or 25°C. We thus established that we could reliably detect survival of *F. tularensis* LVS in fresh water held at 4°C for over two weeks, providing a model system to further interrogate the genetic requirements for environmental persistence.

**Figure 3.**
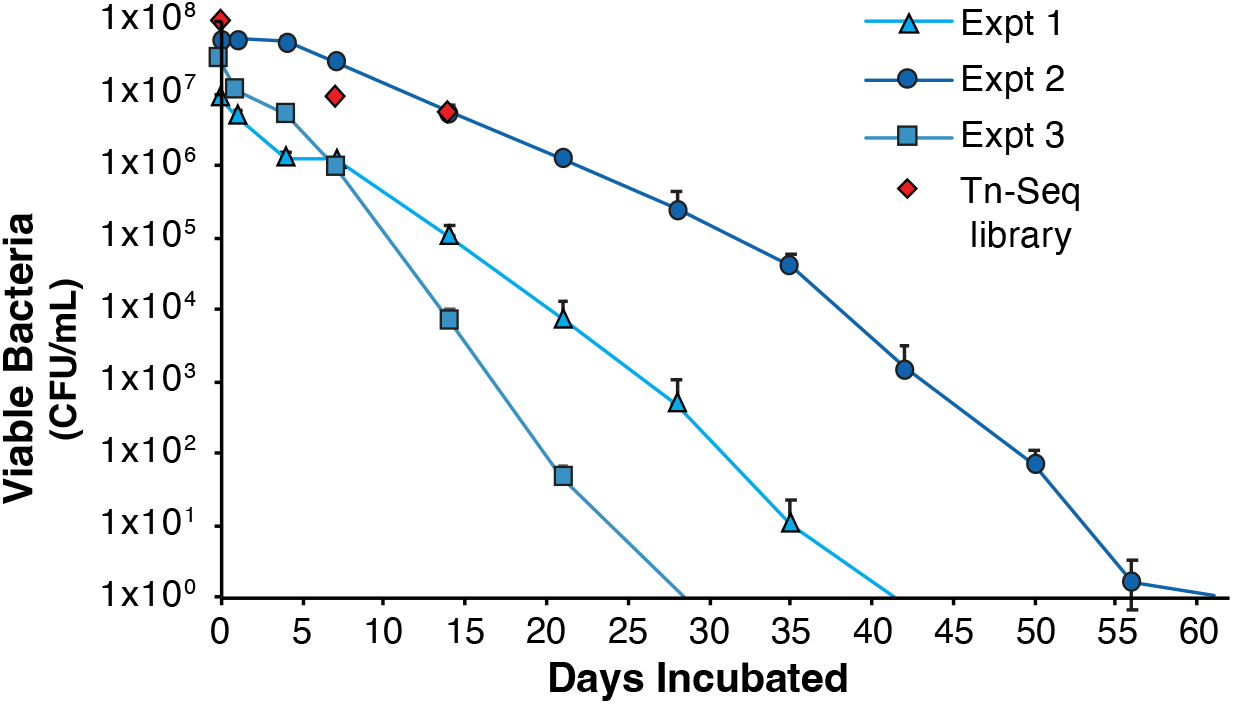
Survival of *F. tularensis* in fresh water held at 4°C. Average CFU per mL recovered at indicated time points for three experiments with cells incubated at 4°C. Experiment 1 data shown is from Figure 2. Viable cells recovered from experiment 1, 2 and 3 are represented by blue triangles, blue circles, and blue squares, respectively. The number of viable cells from the transposon mutant library and when cells were collected for analysis by transposon insertion sequencing (days 0, 7, and 14) are indicated by red diamonds. Each point represents the average number of cells recovered from three independent flasks. Error bars represent 1 SD.

### Modification of a transposon delivery vector

A mariner-based transposon delivery vector compatible with the INSeq protocol for high-throughput sequencing library preparation was previously developed for *F. tularensis* LVS (Ramsey, Ledvina, et al., 2020). However, while sequencing libraries prepared from mutants made with this vector are compatible with older Illumina single-end read flow cells, they are not compatible with newer Illumina chemistry. In particular, sequences corresponding to a shorter Illumina P7 flow site attachment are encoded in the delivery vector (Goodman, 2009). Illumina sequencing kits used with newer instruments, including MiSeq instruments, require a longer P7 sequence and are not compatible with the original, shorter P7 site. Given the relatively small size of the *F. tularensis* genome (∼2 Mb), using a MiSeq instrument to generate data from transposon insertion libraries would be faster and more cost-effective. Thus, we modified the previously-published *F. tularensis* transposon delivery vector (pKL91; Ramsey, Ledvina, et al., 2020), adding the DNA specifying the full-length P7 site to both inverted repeat regions. This resulted in a new *F. tularensis* transposon delivery vector, pKR141.

### Creation of a transposon mutant library and survival of mutants in fresh water

Using our new transposon delivery vector, which encodes kanamycin-resistance within the transposon, we generated a library of *F. tularensis* LVS mutants. Specifically, we mutagenized wild-type cells ten independent times and pooled ∼8,000 independent kanamycin-resistant colonies.

To determine what genes might be necessary for *F. tularensis* to persist in aquatic environments, we used a Tn-Seq strategy to identify which transposon mutants are unable to survive in our established fresh water persistence model. We started by resuspending cells from our mutant library into fresh water. These cells were used to inoculate three independent flasks, which were sampled to assess the initial number of viable bacterial cells, and the remaining sample was saved to identify the mutants present in the original library inoculum. After seven and fourteen days of incubation at 4°C, we sampled the flasks to assess bacterial viability (**Fig. 3**) and to extract genomic DNA to determine which mutants remained. We extracted genomic DNA from (1) cells used as the inoculum, (2) cells remaining after seven days and (3) cells remaining after fourteen days. We used this DNA to specifically amplify the junction between the genomic DNA and transposon insertion and created libraries for high-throughput sequencing (Goodman et al., 2011). While we were able to obtain sequencing data using a MiSeq, inefficient clustering led to us using a relatively low numbers of reads for our analysis (less than 2 million). Regardless, we were able to identify 5,733, 6,821, and 6,168 unique insertions in the inoculum, day 7, and day 14 samples, respectively. Combining these data, we found 7,818 unique insertion sites in our libraries. While these insertions do not represent a large percentage of the overall number of potential insertion sites (4% of the total TA sites in the genome, and 5% of the non-essential TA sites; Ramsey, Ledvina, et al., 2020), it does correspond to almost four insertions for every annotated gene on average.

Comparison of the transposon mutants present in each condition revealed that mutants in one gene in particular, *mpl*, were lost over time (**Fig. 4**). While our experiment began with three insertion mutants in this gene, only two remained after 7 days and none were recovered after 14 days. This suggests that the product encoded by *mpl*, murein peptide ligase, is key for long-term survival of *F. tularensis* LVS in fresh water.

**Figure 4.**
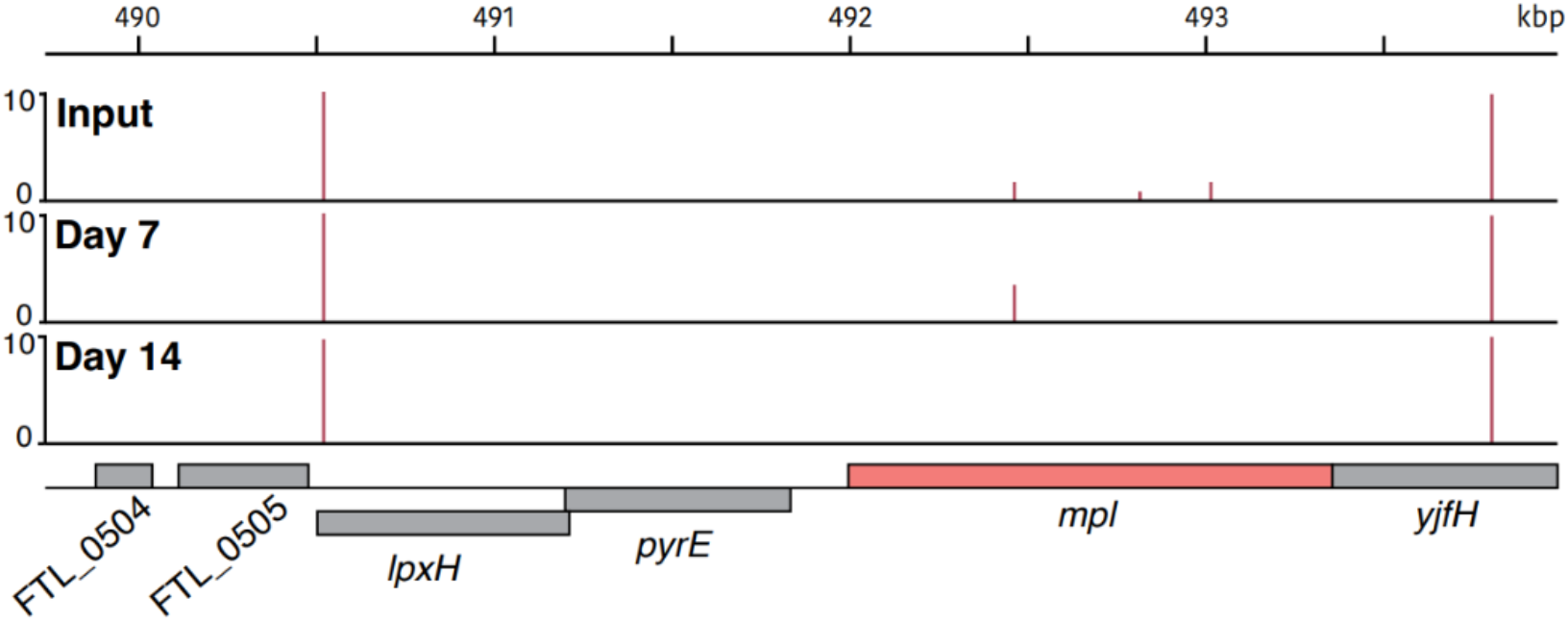
Insertion mutants in the *F. tularensis* LVS *mpl* gene are unable to persist in fresh water held at 4°C. Transposon insertion profiles from the input library, mutants recovered at day 7, and mutants recovered at day 14 in the chromosomal region surrounding *mpl* (colored in salmon). Line height represents the relative abundance of sequencing reads at that position on a log scale.

## Discussion

In this work, we established a laboratory model system to examine the persistence of *F. tularensis* in fresh water using the live vaccine strain (LVS). In establishing this model, we found *F. tularensis* LVS cells could remain viable for 3 – 8 weeks in cold (4°C) fresh water. Using this model and mutants generated using a modified transposon delivery vector, we performed a transposon insertion sequencing assay to identify genes necessary for long-term persistence of *F. tularensis* in fresh water. We found that mutants with insertions in one particular gene, *mpl*, were unable to persist in this model. This suggests that the *mpl* gene product, murein peptide ligase, is important to allow cells to remain viable in this hypoosmotic stress condition.

Given that viable *F. tularensis* cells in the environment are likely major drivers of human infection, a number of studies have examined a variety of conditions that impact persistence of *F. tularensis* (Forsman et al., 2000; Thelaus et al., 2009; Berrada et al., 2011; Gilbert and Rose, 2012; Siebert et al., 2020; Hennebique et al., 2021; Golovliov et al., 2021). The conditions we found most favorable for *F. tularensis* LVS persistence in fresh water, incubation at 4°C, is consistent with many of these prior studies. We did observe variability in the length of time bacteria remained viable at 4°C, with the last viable cells detected between 21 to 56 days. This may be related to some variability in the number of starting cells; our second experiment began with the highest number of viable cells and these cells remained viable for the longest amount of time. It is possible that the larger number of dying cells provided additional nutrition, allowing longer persistence of the remaining cells.

The modifications we made to our transposon delivery vector aided our transposon insertion sequencing screen, allowing us to make and sequence libraries quickly and in a cost-effective manner. However, our identification of genes important for persistence in fresh water is limited by the original size of the transposon mutant library (fewer than 10,000 mutants) which is not highly saturated. We predict that this limitation precluded identification of additional genes required for long-term viability in fresh water. Future experiments would benefit from a larger mutant library and will use optimized library sequencing conditions to provide significantly more read depth.

One gene expected to be important for aquatic survival of *F. tularensis* is FTL_1753, encoding the mechanosensitive channel FtMscS. The presence of FtMscS protects *F. tularensis* during hypoosmotic shock, allowing increased viability (Williamson et al., 2018). Notably, we did not observe large differences in survival of FTL_1753 mutants in our experiment. We hypothesize this is because we compared cells which had already experienced hypoosmotic shock (specifically, cells resuspended in fresh water) to those which remained viable over long periods of time (after 7 or 14 days in fresh water). This suggest that while FtMscS is likely crucial for the transition from host cells into fresh water environments, it is not strictly necessary for long-term viability in cells which have already experienced hypoosmotic shock.

Importantly, our study reveals that mutants in the *F. tularensis mpl* gene are defective for persistence in fresh water. The *mpl* gene product, UDP-N-acetylmuramate:L-alanyl-γ-D-glutamyl-*meso-*diaminopimelate ligase or murein peptide ligase, functions in peptidoglycan recycling (Mengin-Lecreulx et al., 1996; Hervé et al., 2007). In particular, Mpl ligates the peptidoglycan turnover product L-Ala-γ-D-Glu-*meso*-diaminopimelate to UDT-*N*-acetylmuramic acid (UDP-MurNAc) in the cytosol, which can then enter the *de novo* peptidoglycan biosynthesis pathway. While maintenance of an intact cell envelope is crucial for bacterial survival, prior work and this study demonstrate that *mpl* is not essential in *F. tularensis* LVS, nor is it essential in *E. coli* (Ramsey, Ledvina et al., 2020; Mengin-Lecreulx et al., 1996). However, *mpl* was identified as critical for *F. tularensis* survival in macrophage. The conditional requirement for *mpl* may reflect a necessity for peptidoglycan recycling to maintain cell wall integrity during diverse stress conditions (intramacrophage growth and hypoosmotic stress).

A study of genes required for another intracellular pathogen, *Legionella pneumophila*, to survive in fresh water similarly identified a gene necessary for cell wall maintenance, the L, D - transpeptidase *lpg1697* (Aurass et al., 2023). Both Lpg1697 and Mpl function in peptidoglycan remodeling, raising the possibility that this is a required activity for diverse bacteria during the stress encountered in long-term hypoosmotic environments.

## Materials and Methods

### Bacterial strains and growth conditions

Except where otherwise noted, *Francisella tularensis* subsp. *holarctica* LVS were grown aerobically at 37°C on cystine heart agar (Difco) supplemented with 2% hemoglobin (CHAH) or in Mueller Hinton broth (Difco) supplemented with glucose (0.1%), ferric pyrophosphate (0.025%), and Isovitalex (2%). *Escherichia coli* strain PIR-1 (Invitrogen) was used in vector construction. For selection, kanamycin was used at 5 µg/mL (LVS) or carbenicillin was used at 10 µg/mL (*E. coli*).

### Cell Survival Assay

Fresh water was collected from the Beaver River in Rhode Island and sterilized by filtration using a 0.22-micron filter. Wild-type LVS or transposon mutant library cells were grown to a confluent lawn on CHAH and resuspended in sterile water to a final OD_600_ of 0.03. Cells in fresh water were incubated at indicated temperatures (4°C, 16°C, or 25°C) in triplicate. The cultures were serially diluted and plated on CHAH to determine the viable colony forming units (CFU) on indicated days. For the wild-type cells, when viable CFUs dropped to below ∼10^4^, 0.3 mL of undiluted sample were plated for enumeration and, after detecting the last viable CFU, samples were plated for two additional weeks to confirm no viable cells remained. Plates were incubated at 37°C for 3-4 days or until single colonies could be counted. This cell survival assay was completed three times. Cells were removed from transposon mutant library cultures for genomic DNA isolation on the same day they were plated for CFU.

### Vector Construction

The previously used transposon delivery vector, pKL91, was modified for compatibility with the current Illumina P7 capture site, as this site is encoded within the transposon inverted repeats. A two-step process was used to modify both ends of the transposon. First, pKL91 was digested with KpnI and PacI to remove the sequence encompassing the inverted repeat region at one end of the transposon. A dsDNA fragment was synthesized by IDT that contains the sequence between the KpnI and PacI sites, modified only by the addition of the additional nucleotides necessary to encode the entire P7 capture site (5’ CAA GCA GAA GAC GGC ATA CGA GAT 3’). The synthesized fragment was digested by KpnI and PacI and ligated into the digested pKL91 to yield pKR140. This process was repeated to modify the P7 capture site on the other end of the transposon, digesting pKR140 with PstI and BamHI and replacing the DNA with a synthesized fragment containing the full 24 bp P7 site. Both pKR140 and pKR141 were verified by Sanger sequencing (Rhode Island INBRE Molecular Informatics Core).

### Transposon mutant library construction

To generate the transposon mutant library, *F. tularensis* LVS cells were transformed essentially as described (Maier et al., 2004) using one microgram of transposon delivery plasmid pKR141. Ten independent electroporations were performed and transposon mutants were selected by plating on CHAH with 5 µg/mL kanamycin. The kanamycin-resistant colonies were combined and frozen, approximately 8,000 colonies. A single aliquot of the combined transposon mutants was spread on CHAH plate containing 5 µg/mL kanamycin and incubated at 37°C to a confluent lawn. These cells were resuspended in fresh water and used in the initial inoculation of the fresh water media for Tn-Seq experiment.

### INSeq library construction and sequencing

Sequencing libraries were generated from genomic DNA corresponding to three samples: (1) all the transposon mutant cells grown on solid media and resuspended in fresh water, (2) mutant cells present after incubation in fresh water for 7 days at 4°C, and (3) mutant cells present after incubation in fresh water for 14 days at 4°C. Genomic DNA was isolated as described (Ramsey, Ledvina et al., 2020). Sequencing libraries were generated as described (Goodman et al., 2011), using the following modified primers: BioSamA_v2: 5’ – AAG ACG GCA TAC GAG ATT ACG AAG ACC – 3’, LIB_PCR_5_v2: 5’ – CAA GCA GAA GAC GGC ATA CGA GAT TAC GAA GAC CGG GGA CTT ATC ATC CAA CCT GT – 3’. The three libraries were sequenced using two 150 cycle V3 kits (Illumina) with a MiSeq (Illumina; Rhode Island INBRE Molecular Informatics Core).

### Tn-Seq data analysis

Sequencing reads were demultiplexed, trimmed, mapped and insertions in TA sites were tallied as previously described (Ramsey, Ledvina et al., 2020). Examination of these data revealed one gene, FTL_0508 or *mpl*, which harbored multiple insertions at days 0 and 7 but zero insertions at day 14.

## Acknowledgements

We thank the other members of the Ramsey laboratory (particularly Sierra S. Schmidt for helpful discussions and comments on the manuscript), Janet Atoyan, and other staff at the the Rhode Island INBRE Molecular Informatics Core. Funding for this work was provided by the USDA National Institute of Food and Agriculture, Hatch Formula project accession number 1017848. The research was made possible using funding for AM as well as equipment and services available through the Rhode Island Institutional Development Award (IDeA) Network of Biomedical Research Excellence from the National Institute of General Medical Sciences of the National Institutes of Health under grant number P20GM103430 and through the Centralized Research Core facility and the Molecular Informatics Core (RRID:SCR_017685).

